# Genomic reconstruction of *Bacillus anthracis* from complex environmental samples enables high throughput identification and lineage assignment in Pakistan

**DOI:** 10.1101/2025.01.21.634099

**Authors:** Justin C Podowski, Sara Forrester, Tahir Yaqub, Amin Aqel, Mohammad Abu-Lubad, Nadia Mukhtar, Muhammad Waqar Aziz, Nageen Sardar, Hassaan Bin Aslam, Hamda Pervaiz, Alan J Wolfe, Daniel S Schabacker

## Abstract

*Bacillus anthracis*, the causative agent of anthrax, is a highly virulent zoonotic pathogen primarily affecting domesticated and wild herbivores. Human exposure to *B. anthracis* is primarily through contact with infected animals or contaminated animal products. In Pakistan, where livestock vaccines are largely unavailable and infected carcasses are often disposed of improperly, the risk to humans, wildlife and livestock is significant. Currently, diagnosis of anthrax infections and outbreak tracing necessitates the isolation and culturing of *B. anthracis*, a process that requires BSL-3 facilities. In this study, we show that positive identification, genome reconstruction and lineage assignment can be accomplished using bioinformatic analysis of DNA extracted directly from environmental samples that would otherwise provide the starting material for isolation and culturing. This approach does not require laboratory target enrichment as is necessary for other pathogens, due in part to the extremely high bacterial load in the bloodstream in the deceased animals. Using these methods, we greatly expand the knowledge of endemic *B. anthracis* in Pakistan. We provide the first reference *B. anthracis* genomes from Pakistan since the 1970s and identify A.Br.014 Aust94 as a minor circulating sublineage alongside dominant A.Br.047 Vollum. Future work will focus on limits of detection and will determine if this bioinformatic method can be expanded more broadly for *B. anthracis* or other pathogens to replace typical culture-based methods.

## Introduction

*Bacillus anthracis* is a Gram-positive bacillus that causes anthrax, a high mortality infection of humans and many wild and domesticated mammals. Soil is the natural reservoir for *B. anthracis*, and as a result infection in wild and domesticated animals occurs largely through grazing. Infections in humans occurs through contact with animal carcasses, meat and other animal products derived from those animals. As a result, major factors determining the rates of human infections are endemicity of *B. anthracis* in the soil, and the degree of interaction between humans and infected animals or animal products. While *B. anthracis* is present on all continents except Antarctica (Ert et al., 2007), continents with the greatest risk to humans are Africa, Eurasia and North America (Carlson et al., 2019). Risks in Pakistan are particularly high due to low rates of livestock vaccination (Carlson et al., 2019; Sardar et al., 2023b) and regular interaction with livestock due to large rural populations (Ali and Ejaz, 2023), including meat scavenging from carcasses (Sardar et al., 2023a). As a result, *B. anthracis* is found routinely in soils inside of villages (Rashid et al., 2018), suggesting the potential for non-zoonotic infection of humans directly from soil. A more complete understanding of the prevailing lineages of *B. anthracis* in Pakistan is vital to successful outbreak tracing, but very few sequenced isolates exist (Abdel-Glil et al., 2021; Bruce et al., 2020), limiting these efforts.

Detection of *B. anthracis* in a clinical or environmental sample can be accomplished through amplification of key infectivity genes (Shabbir et al., 2015). However, this does not allow for the phylogenetic inference of lineages, which is necessary for outbreak tracing and usually requires isolation of *B. anthracis* through culturing, DNA extraction and either variable number tandem repeat (VNTR) analysis (Ert et al., 2007) or bioinformatic analysis (Abdel-Glil et al., 2021). The limited availability of BSL-3 facilities necessary to culture *B. anthracis* in many countries most threatened by *B. anthracis* limits outbreak tracing (Peters, 2019). Culture-independent analysis of other pathogens (Flurin et al., 2022) suggests that environmental samples could be used to identify and reconstruct *B. anthracis* genomes.

Compared to other pathogens, the population structure and infection dynamics of *B. anthracis* present both challenges and opportunities to culture independent analysis. High similarity between *B. anthracis* and related *Bacillus cereus* group organisms complicates the use of small-subunit (SSU) based marker approaches (Ehling-Schulz et al., 2019; Helgason et al., 2000), and cryptic presence of infectivity plasmids in *Bacillus cereus* group organisms complicates use of these infectivity genes (Okinaka et al., 2006). However, extremely high abundance of *B. anthracis* in the blood of infected animals (Klein et al., 1966) means that amplification or target enrichment may not be necessary, as it is for other pathogens (Peng et al., 2024).

Here we show that *B. anthracis* complete genomes can be reconstructed from blood-stained soil samples collected near animals that had died from anthrax infections, and that these recovered genomes can be used to identify lineages necessary for outbreak tracing. Overall, we demonstrate the feasibility of culture independent methods to surveil *B. anthracis*, while broadly expanding knowledge of endemic *B. anthracis* in Pakistan.

## Methods

### Sample Collection

Between March 2019 and July 2021, a total of 570 soil samples were collected as described previously (Sardar et al., 2023a). In brief, an outbreak was identified if dead livestock exhibited signs consistent with anthrax (blood oozing from carcass’s orifices, bloating, absence of rigor mortis). Blood-stained soil samples were collected near the mouth or anus of the animal. Several samples per outbreak were collected. Blood-stained soil samples were processed for the isolation of *B. anthracis* spores using the previously described GABRI method (Fasanella et al., 2013). Samples processed in this work were restricted to those that were Gram-positive, containing green-colored spores in elongated chains, and were PCR-positive for *cap, lef*, and *ef* genes (Fasanella et al., 2003). DNA was extracted from samples using a QIAamp DNA Mini Kit (Qiagen, Germantown, MD) for metagenomic analysis. For *B. anthracis* isolates, blood-stained soil was used to generate cultures using selective media (PLET Agar) and differential media (Blood Agar). DNA was extracted from isolates using QIAamp DNA Mini Kit (Qiagen, Germantown, MD). All DNA samples were subjected to Health and Human Services (HHS) regulations to ensure no viable *B. anthracis* was present in DNA extracts (HHS validation of BA inactivation, HHS-0920-2018-F-2274), as well as USDA regulations to ensure no viable Foot-and-mouth disease virus was present. DNA samples were then shipped to the United States for sequencing.

### DNA Sequencing

For all samples, library preparation was carried out using the Illumina DNA Prep tagmentation kit (Illumina, San Diego, CA). Isolated genome samples were sequenced on an Illumina NextSeq 2000 at 400Mbp at Microbial Genome Sequencing Center LLC (Pittsburgh, PA). Complex samples were sequenced on an Illumina NextSeq 2000 at 2Gbp. For selected complex samples ZBM8S2_S1, ZBM8S3_S2, ZBM8S5_S3, ZBM8S6_S4, ZBM11S2_S5, ZBM11S4_S6, ZBM11S7_S7 library preparation and sequencing was instead carried out at Argonne National Laboratory Environmental Sample Preparation and Sequencing Facility. Library preparation was carried out using PrepX™ DNA Library Kit (Takara Bio USA, San Jose, CA) and samples were sequenced on an Illumina NextSeq2000 at 45 Gbp.

### Bioinformatic Analysis

Unless otherwise stated, defaults were used for bioinformatic software. Raw reads from isolate *B. anthracis* genomic data were quality controlled using BBTools v39.06 (Bushnell et al., 2017). BBduk with ktrim=r k=23 mink=11 hdist=1 tpe tbo was used to remove adapters, removehuman.sh was used to remove human reads, BBduk trimq=30 qtrim=r was used to quality control, and BBnorm target=100 mindepth=2 was used to normalize read depth. SPAdes v3.15.5 (Bankevich et al., 2012) was used to assemble these reads into contigs with the –isolate flag. Stats.sh from BBTools was used to assess contig size.

Raw reads from complex samples had adapters removed, human reads removed, and sequences quality controlled using same methods as for isolate genome samples. To assess the percentage of a short read sample that was definitively *B. anthracis*, krakenuniq (Breitwieser et al., 2018) was used to compare sequences against the precompiled MicrobialDB, for both whole genome samples and complex samples. Multiple distinct methods were then used to test the efficiency of extraction of *B. anthracis* DNA from complex samples.

Multiple bioinformatic methods (untargeted, semi-targeted, targeted, highly targeted) were tested to determine how *B. anthracis* genomes could be most efficiently recovered. Untargeted and semi-targeted methods that used metagenomic binning approaches were attractive because they allow for isolation without a reference, which could potentially enforce results similar to reference used. Whereas targeted and highly targeted methods have the potential downside of enforcing similar results to the reference used, they are much quicker and required fewer computational resources.

For method 1 (untargeted), reads were normalized using BBnorm target=100 mindepth=2 and then assembled using megahit v1.2.9 (Li et al., 2015) with –presets meta-sensitive. Metagenomic binning was carried out in metaWRAP v1.3.2 (Uritskiy et al., 2018). Bowtie2 v2.3.5.1 (Langmead and Salzberg, 2012) was used to generate coverage for contigs and metabat2 v 2.12.1 (Kang et al., 2019), maxbin2 v2.2.6 (Wu et al., 2016) and CONCOCT v1.0.0 (Alneberg et al., 2014) were all used to identify bins. MetaWRAP bin_refinement was used with -c 75 -x 5 to identify high completions bins using CheckM v1.0.12 (Parks et al., 2015).

For method 2 (semi-targeted), reads were normalized and assembled as in method 1. Assemblies were then submitted to the Bacterial and Viral Bioinformatic Resource Center (BV-BRC) (Olson et al., 2023) where metagenomic binning was carried out (Parrello et al., 2021) using what we deemed ‘semi-targeted’ binning.

For method 3 (targeted), a bowtie2 index was created using the *B. anthracis* ‘Ames Ancestor’ strain (GCF_000008445.1) and reads were mapped against that index using bowtie2 --sensitive --no-unal. Read pairs where both reads were mapped were extracted from the resultant samfile, and SPAdes was used to assemble using –isolate.

For method 4 (highly targeted), analysis was identical to method 3 except --trusted-contigs to the GCF_000008445.1 assembly fasta was used. This was done to enforce reference-based assembly.

GTDB-Tk v2.1.1 (Chaumeil et al., 2022) with database release 214 was then used to identify taxonomy of genomes from all methods to identify *B. anthracis*. Once high-quality *B. anthracis* genomes were identified from either isolates or complex samples, CanSNPer v1.0.10 was used to assign sublineage, and PubMLST (Jolley et al., 2018) was used to assign cgMLST profiles specific to *B. anthracis* (Abdel-Glil et al., 2021).

ArcGIS Pro v3.2 was used to generate maps depicting geographic distribution of sampling and cgMLST assignments.

## Results

Samples were collected from 13 outbreaks across the districts of Zhob, Bajour, and Bahawalnagar in Pakistan. Genomic DNA from 10 isolates was collected across 5 outbreaks and paired with the metagenomic DNA from the complex sample (blood-stained soil) that generated each of these 10 isolates. An additional 21 complex samples were also collected to expand surveillance of sublineages beyond outbreaks where isolates could be generated.

### Isolate Genomes

High quality isolate genomes were successfully obtained from all 10 isolate samples. For all 10 samples, krakenuniq identified 99% of the reads in each sample as *Bacillus cereus* group and identified at least 78%-81% of the reads specifically as *B. anthracis*. Assembled genome sizes ranged from 5.5 Mb to 6.5 Mb and, on average, 93% of those genomes were contained within contigs larger than 50Kb (Table 1).

**Table 1:** Description of the 10 sequenced isolate genomes.

| Sample Name | cgMLST | % reads *B. anthracis* | Isolate genome size | % genome in scaffolds > 50kb |
| --- | --- | --- | --- | --- |
| ZBMS4 | A.Br.047 | 79% | 5.470 MB | 96% |
| ZBMS6 | A.Br.047 | 79% | 5.476 MB | 96% |
| ZBM2S3 | A.Br.047 | 78% | 5.471 MB | 96% |
| ZBM2S5 | A.Br.047 | 78% | 5.501 MB | 95% |
| ZBM11S7 | A.Br.047 | 78% | 5.473 MB | 96% |
| ZBM11S8 | A.Br.014 | 79% | 5.727 MB | 92% |
| BW7S4 | A.Br.014 | 79% | 6.490 MB | 81% |
| BW7S5 | A.Br.014 | 79% | 5.468 MB | 96% |
| BW10S3 | A.Br.014 | 80% | 6.251 MB | 82% |
| BW10S5 | A.Br.014 | 80% | 5.472 MB | 95% |

5 isolate genomes were assigned to the A.Br.007 Vollum clade, and 5 were assigned to the A.Br.003 Aust94 clade, using CanSNPer (Ert et al., 2007; Lärkeryd et al., 2014). Concordantly, the 5 assigned A.Br.007 were identified more specifically as A.Br.047 by cgMLST (Abdel-Glil et al., 2021; Jolley et al., 2018), while those assigned A.Br.003 were more specifically assigned A.Br.014. The closest existing isolate for the A.Br.047 genomes was SK-102 (GCF_000832565.1), which was isolated from wool originating in Pakistan in 1976, and the closest isolate for A.Br.014 genomes was A0656 (GCA_029700625.1), which was isolated from soil in China in 1982.

### Genome Reconstruction

Thirty-one DNA samples derived from blood-stained soil were sequenced. For these 31 samples, the percentage of positively identified *B. anthracis* sequences ranged from 1.65% to 81%. Of these 31 samples, 3 samples did not produce a reconstructed *B. anthracis* genome using any method. Two of these 3 samples produced reconstructed genomes identified as *B. cereus*, while 1 produced a reads sample identified as *B. anthracis*, an average of 6% compared to the overall average of 46% across all samples. However, some samples with a lower percentage of reads identified as *B. anthracis* did produce reconstructed *B. anthracis* genomes.

Across 4 bioinformatic methods – untargeted, semi-targeted, targeted and highly-targeted-we attempted to reconstruct *B. anthracis* genomes from which lineage information could be assigned (Table 2). Untargeted methods produced a *B. anthracis* genome in 20 samples, semi-targeted in 23 samples, targeted in 27 samples and highly targeted in 31 samples (Table S1). However, our highly targeted method generated false positives, with 3 cases displaying results consistent with the *B. anthracis* ‘Ames Ancestor’ strain used as reference in the highly targeted assembly, in the samples where untargeted methods indicated another *B. cereus* group organism was present in that sample. Across all samples, cgMLST and CanSNPer results were always consistent within a sample.

**Table 2:** Description of the four bioinformatic methods used.

|  | Untargeted Method 1 | Semitargeted Method 2 | Targeted Method 3 | Highly Targeted Method 4 |
| --- | --- | --- | --- | --- |
| Genomic content targeting | After Assembly | After Assembly | Before Assembly | Before Assembly |
| Genomic content targeting method | MetaWrap - CONCOCT, metabat2, maxbin2 | BV-BRC metagenomic binning | Bowtie2 read recruitment | Bowtie2 read recruitment |
| Assembly method | Megahit | Megahit | SPAdes | SPAdes reference assembly |
| Taxonomic assignment | GTDBtk | GTDBtk | GTDBtk | GTDBtk |
| CanSNP assignment | CanSNPer *Bacillus anthracis* | CanSNPer *Bacillus anthracis* | CanSNPer *Bacillus anthracis* | CanSNPer *Bacillus anthracis* |
| cgMLST assignment | PubMLST *B. anthracis*<br>cgMLST | PubMLST *B. anthracis*<br>cgMLST | PubMLST *B. anthracis*<br>cgMLST | PubMLST *B. anthracis*<br>cgMLST |

In 10 cases where isolates had been obtained, genomes were generated from both the complex samples and the resulting isolates. Of these 10 cases, 7 complex samples produced reconstructed *B. anthracis* genomes with sublineage assignments that matched the sub lineage assignment of the isolate genome. For the 3 cases that did not match, all were cases in which A.Br.047 was assigned to the reconstructed *B. anthracis* from the complex sample while A.Br.014 was assigned to the isolate *B. anthracis*. In two of the three cases, A.Br.047 was also present in other samples, including other isolates, in the same outbreak. This may suggest that the disagreement between isolate and complex sample assignments could be due to multiple sublineages present in a sample. In one case where A.Br.047 was not otherwise detected, we sequenced a second extract of the complex sample and recovered a A.Br.047 assignment the second time.

Samples were collected across 13 outbreaks in Zhob, Bajour and Fortabbas in Pakistan. Each outbreak contained a single dead animal, and each sample came from blood-stained soil surrounding that animal. 17 samples were collected from soil surrounding sheep, 3 from goats and 11 from cows. We assigned sublineage to at least one sample across all 13 outbreaks. An outbreak was defined as a single identified animal, and as a result all samples from an outbreak came from blood-stained soil near a single animal carcass. Sublineage A.Br.047 Vollum was by far the most dominant sublineage and was present in all 13 outbreaks (Figure 1, Table 3). Sublineage A.Br.014 Aust94 was found in 2 of 3 outbreaks sampled in Bajour but only 1 of 9 outbreaks in Zhob.

**Table 3:** Samples summarized by outbreak, including cgMLST assignments.

| A.Br.014 | A.Br.047 | Outbreak | Province | Country | Animal | Outbreak Date |
| --- | --- | --- | --- | --- | --- | --- |
| 0% | 100% | 5 | Zhob | Pakistan | Goat | Apr-2019 |
| 0% | 100% | 44 | Fortabbas | Pakistan | Cow | Mar-2021 |
| 0% | 100% | 40 | Zhob | Pakistan | Sheep | Jul-2020 |
| 33.3% | 66.6% | 37 | Zhob | Pakistan | Sheep | Jun-2020 |
| 60% | 40% | 36 | Bajour | Pakistan | Cow | Oct-2019 |
| 0% | 100% | 35 | Zhob | Pakistan | Sheep | Oct-2019 |
| 0% | 100% | 33 | Zhob | Pakistan | Sheep | Sep-2019 |
| 50% | 50% | 29 | Bajour | Pakistan | Cow | Sep-2019 |
| 0% | 100% | 26 | Zhob | Pakistan | Sheep | Aug-2019 |
| 0% | 100% | 24 | Zhob | Pakistan | Sheep | Aug-2019 |
| 0% | 100% | 17 | Zhob | Pakistan | Sheep | Aug-2019 |
| 0% | 100% | 14 | Zhob | Pakistan | Cow | Jun-2019 |
| 0% | 100% | 13 | Bajour | Pakistan | Cow | Jul-2019 |

**Figure 1.**
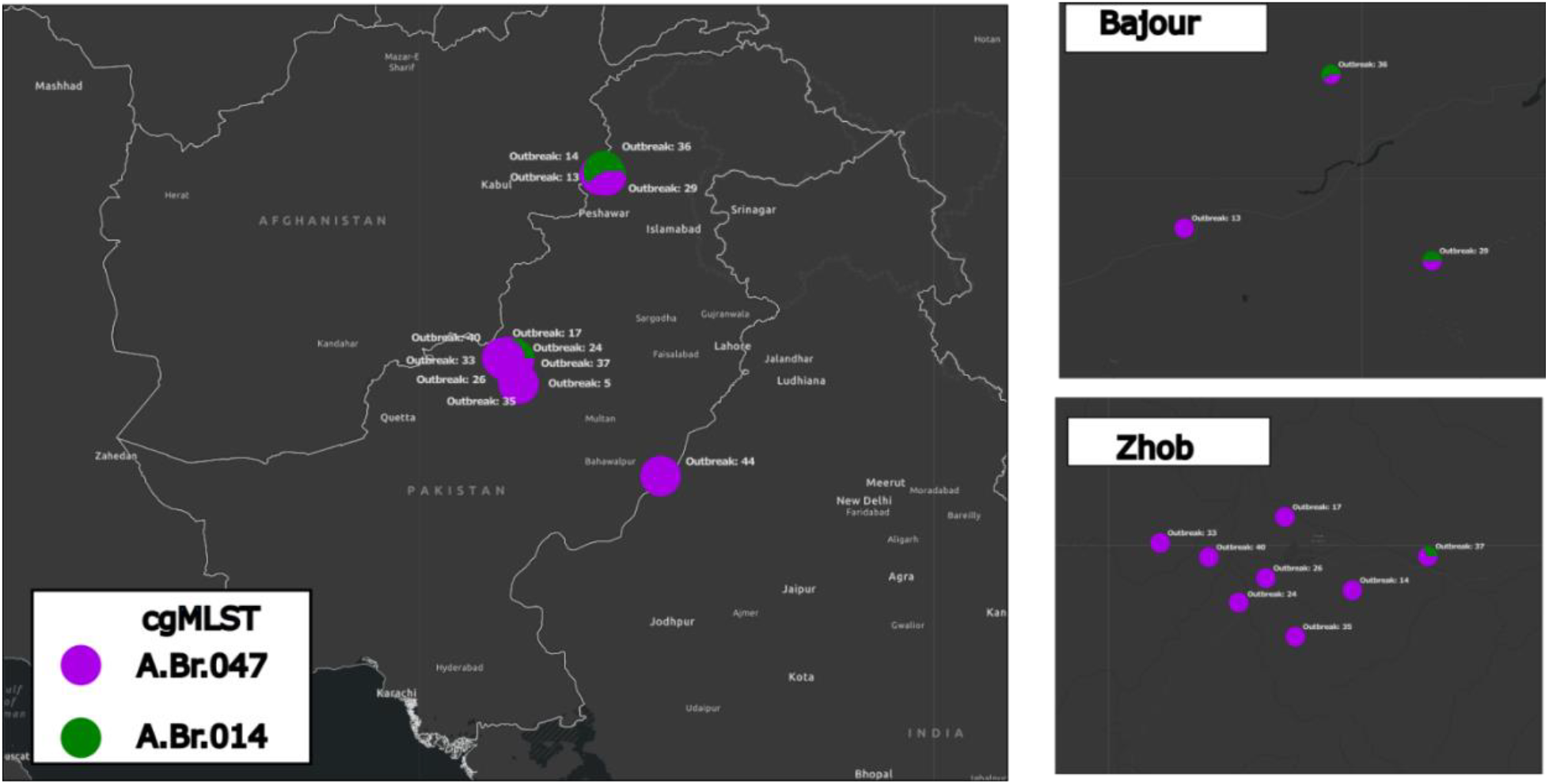
GIS Map of outbreaks in Pakistan, with pie charts colored by proportion of cgMLST assignments for samples in that outbreak. Bajour (northern-most region plotted on whole map) and Zhob (central region plotted on whole map) are districts in Pakistan.

## Discussion

Our results suggest that computational reconstruction of *B. anthracis* can be employed in the absence of the ability to isolate *B. anthracis* and used to assign sublineage through cgMLST. Due in part to high clonality, assembly of *B. anthracis* in complex samples led to reconstructed genomes of sufficient contiguity and with sufficient cgMLST loci to be considered adequate for cgMLST typing (Abdel-Glil et al., 2021). This does not require targeted enrichment of *B. anthracis* and can operate reliably when *B. anthracis* represents at or above 5% of the total sample. Due to the extremely high load of *B. anthracis* in the blood stream of infected animals (Klein et al., 1966), when collecting samples that are mixtures of blood and soil, the percentage of the sample that is *B. anthracis* is on average nearly 50%.

When comparing bioinformatic methods, a targeted approach (method 3) appeared to be optimal, as it produced the highest number of reconstructed genomes without any false positives. Use of reference-based assembly in the highly targeted approach (method 4) resulted in false positives where sufficient *B. anthracis* sequences were not present. This is likely due to the algorithm inserting reference genome DNA sequences when no sequences are provided. Consistency in cgMLST assignments within a sample was compelling, especially as assembly methods varied between megahit (Li et al., 2015) in untargeted and semi-targeted and SPAdes (Bankevich et al., 2012) in targeted and highly targeted methods. While complex sample assignments were not always consistent with isolate genome assignments, it is likely this is due to multiple lineages of *B. anthracis* within a single animal.

We identified multiple cases in which both A.Br.047 Vollum and A.Br.014 Aust94 were identified in samples from the same animal. As this occurred in two isolate genomes from outbreak 37 in Zhob, this cannot be attributed to computational methods. We also saw this in outbreak 36, where both isolate genomes were A.Br.014 Aust94 and the accompanying complex samples were A.Br.047 Vollum and A.Br.014 Aust94, respectively. This suggests that multiple distinct infection events could have occurred in these animals, with different sublineages in each event. Given the high seroprevalence of antibodies targeting *B. anthracis* in otherwise healthy, unvaccinated animals in Pakistan (Sardar et al., 2023b), repeated infections with *B. anthracis* may not be uncommon. Multi-sublineage infections may be detectable in a single complex sample given future methodological improvement. Long read sequencing is likely superior to short read sequencing for detection of these cases, given the ability to physically associate SNPs across longer reads. However higher input DNA requirements for long read sequencing is challenging, considering the harsh treatment requirements necessary for U.S. HHS verification of no viable *B. anthracis* spores. However, long read sequencing using an Oxford Nanopore Minion within Pakistan is a viable alternative.

Our findings that A.Br.047 Vollum is the dominant sublineage in Pakistan is consistent with past surveys (Abdel-Glil et al., 2021; Ert et al., 2007; Sahl et al., 2016), as the majority of *B. anthracis* genomes isolated from Pakistan have been identified as A.Br.047 Vollum. While A.Br.014 Aust94 has been identified in neighboring India (Ert et al., 2007; Sahl et al., 2016), it has not been identified previously in Pakistan. As none of the provinces in which A.Br.014 Aust94 were detected bordered India, it is likely that A.Br.014 Aust94 is endemic throughout Pakistan but has not been detected due to a lack of sampling, rather than recent migration from India.

This study demonstrates that shotgun sequencing of complex samples is a viable method for identifying *B. anthracis* sublineage in the pursuit of outbreak tracking. We do not constrain how quickly a deceased animal needs to be sampled after death, the ratio of blood to soil needed for recovery of *B. anthracis*, or other parameters which will determine the overall usefulness of this method. However, this study represents a typical sampling campaign with difficult to access samples often in remote locations, without cold chain and with sequencing performed out of the country. As a result, our success suggests the general utility of this method.

## Data Availability

Raw reads and assemblies for isolate genomes are available in BioProject PRJNA1159740. Raw reads for complex samples are available in BioProject PRJNA1159740. Documentation of code involved in genome assembly, as well as complex sample analysis, is available at https://github.com/jpod1010/pathogencomplexsamples/blob/main/pipeline_code.

## Acknowledgment

We thank Scott Schlueter for help with generating ArcGIS maps, and James Davis for computational resources and for his help reviewing manuscript drafts. Funding for this work was provided by the Defense Threat Reduction Agency Cooperative Threat Reduction Program (HDTRA11710050) and by the Argonne National Laboratory Directed Research and Development Program (LDRD 2024-0231)

